# Microscopic Phage Adsorption Assay: High-throughput quantification of virus particle attachment to host bacterial cells

**DOI:** 10.1101/2024.10.09.617072

**Authors:** Jyot D. Antani, Timothy Ward, Thierry Emonet, Paul E. Turner

**Affiliations:** Department of Ecology and Evolutionary Biology, Yale University, New Haven, CT 06520, USA; Center for Phage Biology & Therapy, Yale University, New Haven, CT 06520, USA; Quantitative Biology Institute, Yale University, New Haven, CT 06520, USA; Department of Molecular, Cellular and Developmental Biology, Yale University, New Haven, CT 06520, USA; Department of Physics, Yale University, New Haven, CT 06520, USA; Program in Microbiology, Yale School of Medicine, New Haven, CT 06520, USA

**Keywords:** bacteriophage, phage-bacteria interactions, fluorescence microscopy, adsorption

## Abstract

Phages, viruses of bacteria, play a pivotal role in Earth’s biosphere and hold great promise as therapeutic and diagnostic tools in combating infectious diseases. Attachment of phages to bacterial cells is a crucial initial step of the interaction. The classic assay to quantify the dynamics of phage attachment involves co-culturing and enumeration of bacteria and phages, which is laborious, lengthy, hence low-throughput, and only provides ensemble estimates of model-based adsorption rate constants. Here, we utilized fluorescence microscopy and particle tracking to obtain trajectories of individual virus particles interacting with cells. The trajectory durations quantified the heterogeneity in dwell time, the time that each phage spends interacting with a bacterium. The average dwell time strongly correlated with the classically-measured adsorption rate constant. We successfully applied this technique to quantify host-attachment dynamics of several phages including those targeting key bacterial pathogens. This approach should benefit the field of phage biology by providing highly quantitative, model-free readouts at single-virus resolution, helping to uncover single-virus phenomena missed by traditional measurements. Owing to significant reduction in manual effort, our method should enable rapid, high-throughput screening of a phage library against a target bacterial strain for applications such as therapy or diagnosis.

## Introduction

Bacteria and their viruses, bacteriophages (phages), are estimated to be the two most abundant biological groups on Earth (1). Hence, phage infections of host bacterial cells are among the most numerous biological interactions occurring on the planet, with consequences ranging from ecosystem functions such as geochemical cycles to community dynamics occurring within microbiomes of multicellular organisms (2, 3). Beyond the fundamental need to better understand these frequent biological interactions, there is resurgent interest in using phage biotechnology to address human problems (4, 5). For example, phage therapy can be used as an alternative means of controlling bacterial infections due to the concerning rise in antimicrobial resistance (6, 7). Thus, it becomes crucial to gain a comprehensive understanding of phage interactions with bacteria. Classic genetic and biochemical assays have contributed significantly to probing these interactions in key phage-bacteria models over the last century (2, 8, 9). However, a more detailed understanding of phage infections, especially at the level of individual virus interactions with host cells, is desirable (10, 11).

The first steps in any phage-bacterium interaction are encounter, attachment, and binding of a phage particle to a host cell (12, 13). Phages require specific surface-exposed host structures (receptors) to initiate infection. The receptor(s) can include polysaccharides, outer membrane porins and other transporter proteins, cell-wall teichoic acids, or cellular appendages such as flagella and pili (14, 15). Traditionally, studies of phage attachment (termed adsorption) are quantified through assays utilizing mixtures of roughly millions or more phage particles and bacterial cells: phages are mixed with bacteria and the numbers of un-adsorbed (free) particles are experimentally estimated over time to calculate an adsorption rate constant under an assumed model of first-order association between bacteria and phages (13, 16–21). More complicated models have been proposed to explain the observed decline in numbers of free phages: for instance, phages are known to have non-specific (reversible binding) and specific (irreversible binding) interactions with the bacterial cell surface; some models incorporate this information while interpreting the experimental observations from the classical adsorption assays (22). Nevertheless, all these approaches obtain ensemble estimates of the adsorption rate constant across the phage population, without probing possible heterogeneity in the adsorption dynamics between individual virions and bacterial cells.

In reality, phage-bacteria interactions are highly variable (23). Because of such variability, the dwell time – time that a virus particle spends in contact with a single cell – is likely distributed over a wide range. Being able to routinely probe the distribution of dwell time for different types of phages and cells would be valuable for characterizing the heterogeneity and the underlying stochasticity in dynamics of phage adsorption (23, 24).

In the current study, we develop and validate a Microscopic Phage Adsorption (MPA) assay as a method to quantify phage adsorption dynamics at single-virus resolution. We combined widefield fluorescence microscopy to visualize individual phage particles interacting with immobilized bacterial cells. We used a lysine-specific fluorescent dye to label phage particles, eliminating the requirement to genetically engineer a phage-such as to introduce a foreign gene (e.g., fluorescent protein marker), which would represent severe limitations for non-model species where bacterial and phage genetic engineering may not be trivial. We show that our dye-labeling technique is generalizable to many phages which have different morphologies and which infect various bacterial species.

Specifically, we examined three interrelated hypotheses concerning quantification of phage particle attachment to cells. First, we tested the hypothesis that measuring the distribution of viral particle dwell times using standard and generally available fluorescent microscopy techniques could distinguish the differing attachment abilities of individual phages when challenged to infect various host genotypes that differed in cell-surface receptors. This was done by employing widefield fluorescence microscopy and MATLAB-based single particle tracking: we obtained trajectories of individual phages interacting with bacterial cells immobilized on a glass coverslip. We quantified the distributions of trajectory durations, which are readouts of dwell times, for thousands of single virions interacting with bacterial surfaces. Comparisons of the distributions for phage T4 attachment to various *Escherichia coli* genotypes allowed us to test whether virus adsorption differed according to expected effects of cell-surface receptor presentation. Second, using correlation analysis, we tested the hypothesis that the outcomes from our microscopic measurements (MPA assay) agreed with the classical adsorption-assay method when examined via the phage T4 and *E. coli* bacteria model. Third, we tested whether the fluorescent labeling approach was generalizable across coliphages representing different virus taxonomic groups, and whether the MPA assay was broadly useful for studying adsorption of two phages that infected pathogenic *Shigella flexneri* and *Pseudomonas aeruginosa* host bacteria.

Our approach provides a powerful and general means to quantify phage adsorption at the single-virus level, in a high-throughput manner. In contrast to the laborious and low-throughput classical adsorption assays which only provide ensemble estimates, our approach enables efficient quantification of individual virus attachment to bacterial cells. The technique’s versatility affords great potential to test hypotheses concerning phage biology, as well as to aid development of phage-based technologies.

## Results

### Widefield fluorescence microscopy can be used for probing individual phage T4 interactions with host cells

Aiming to develop a generalizable phage labeling protocol, we used an amine-reactive dye that conjugates with all lysine residues, amine-modified oligonucleotides, and other amine-containing biomolecules exposed on any biological surface. The labeling protocol is described in Materials and Methods. First, we confirmed that labeled phages retained their infectivity of their bacterial host cells: we performed standard plaque assays (see Materials and Methods) to confirm that the labeled phages formed plaques on top agar with bacteria (**Fig S1**).

We imaged fluorescently labeled phages using an inverted widefield fluorescence microscope equipped with a 100×, 1.40 NA objective and a scientific CMOS camera (see Materials and Methods for details on the microscopy setup). Labeled phages formed bright foci when excited with epifluorescence light source. We recorded time-lapse movies of the moving foci at 30 frames per second and used custom MATLAB codes to detect and connect the trajectories of individual foci. Sample trajectories of individual T4 phages are shown in **Fig 1**A. From phage trajectories measured in the absence of bacteria, we calculated mean-squared displacement, *MSD*, as a function of characteristic time *τ*, to determine the diffusion coefficients (least-square fits, *MSD* = 4*Dτ*^*α*^; *D* = 3.3 ± 0.02 and *α* = 1 ± 0.01) of the foci (**Fig 1**B). These values agree with the diffusion coefficients of individual T4 phage particles reported in the literature, as measured through dynamic light scattering (25) as well as microscopic analyses(26, 27).

**Fig 1.**
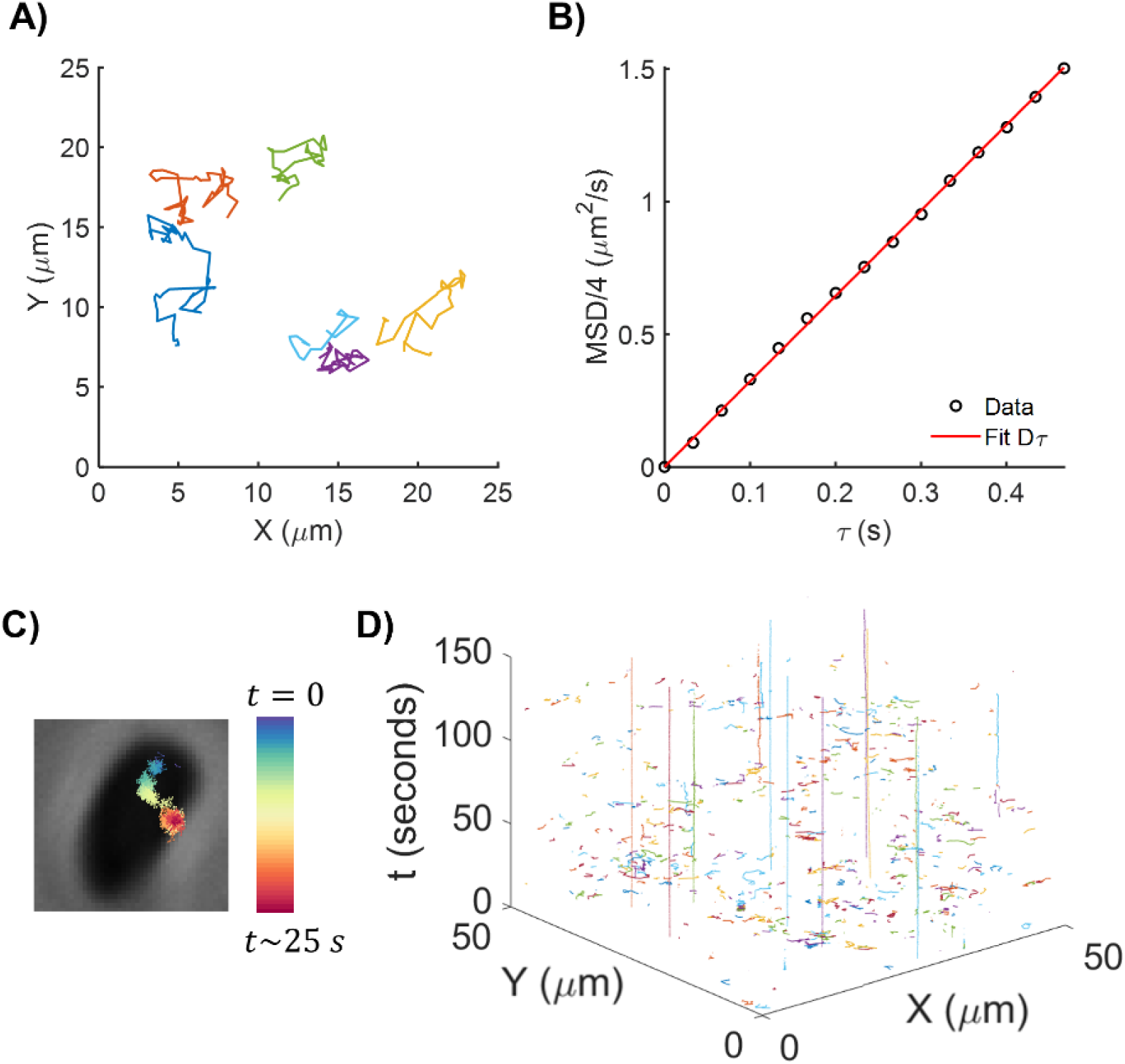
Microscopy-based method to determine trajectories of individual phage particles. **(A)** Trajectories of fluorescently labeled, freely diffusing phages were obtained 10 μm away from glass coverslip. Six examples of individual trajectories are shown. **(B)** Mean square deviation (*MSD*) as a function of characteristic time was calculated from the trajectories (n = 1349 trajectories). Least-squares fit for *MSD* = 4*Dτ*^*α*^ yielded *D* = 3.3 ± 0.2 *μm*^2^/*s* and *α* = 1.00 ± 0.03, confirming that the trajectrories correspond to individual phages. **(C)** For the Microscopic Phage Adsorption assay (MPA assay), phages were introduced to bacteria immobilized on a glass coverslip via poly-L-lysine. Example trajectory of a phage interacting with the underlying bacterial cell for ∼25 s is shown. **(D)** Each experiment yielded thousands of single-phage trajectories. A subset of the trajectories from one experiment is shown with the X-Y coordinates of phages with time as the third axis. A majority of the trajectories are short, representing phages that enter the microscopic field of view for a short time before leaving. Phages that attach to a bacterial cell result in trajectories that remain in the same X-Y neighborhood for an extended time-period, i.e., the long, vertical trajectories on this plot.

Next, we performed high-resolution time-lapse microscopy of phages interacting with bacterial cells. Since temperature is a known factor affecting the adsorption rate (17), we performed all the phage-bacteria experiments at a controlled temperature of 34 °C, the highest temperature stably maintained by our imaging chamber, and close to the 37 °C typical culture conditions. We created a lawn of bacterial cells on glass coverslips coated with poly-L-lysine to immobilize the cells (snapshot included in **Fig S2**). Next, we introduced labeled phages to the sample and started video acquisition. We controlled the concentration of phages such that individual particles were distinguishable and detectable with our MATLAB code. We focused away from the coverslip but still on the bacteria to avoid recording labeled phages that adhered to the poly-L-lysine-coated coverslip. We recorded snapshots of immobilized bacteria (phase-contrast channel) and 2-minute videos of phages (fluorescence channel). We refer to this microscopy protocol as an MPA (Microscopic Phage Adsorption) assay. Performing particle-tracking analysis on the fluorescence videos yielded trajectories of individual phages interacting with bacteria. An example of individual phage trajectory overlaid on underlying bacterial cell is shown in **Fig 1**C.

The focal area in our microscopy was approximately 120 μm × 120 μm, spanning thousands of immobilized bacterial cells packed closely together on the coverslip (**Fig S2**). An example of phage trajectories recorded within a sub-region (50 μm × 50 μm) during the first 150 seconds of acquisition is displayed in **Fig 1**D. Since we track every phage within the focal view, our analysis consisted of particles that were permanently attached, temporarily interacting, as well as potentially non-interacting with bacterial cells. We interpreted the durations of the trajectories according to the phage behavior: non-interacting phages came into the focus of the microscope objective and quickly left within a few frames of recording, resulting in extremely short trajectories – trajectories shorter than 0.1 s were removed from analysis. Phages that temporarily interacted with cells resulted in longer trajectories. Phages that permanently attached were detected at the same position for an extended time: these particles yielded trajectories that were longest in time-axis (i.e., long vertical lines in **Fig 1**D). With relatively close packing of bacteria (**Fig S2**), we assumed that each trajectory longer than 0.1 s likely interacts with a bacterial cell; thus, the duration of a trajectory represents a readout of the dwell time.

To confirm the viability of fluorescently labeled phages at a microscopic level, we performed an experiment where we visualized phages attached to bacterial cells (in the fluorescence channel) and recorded time-lapse videos of the bacteria in the phase-contrast channel for an extended period. A majority of the cells in the focal region of the microscope objective eventually lysed, confirming lytic infections (Movie S1). The multiplicity of infection (MOI) in our experiments was ∼0.01; hence, we ruled out lysis from without (which occurs at a high MOI (28)) as a mechanism of the observed lysis. Finally, we confirmed through classic adsorption assays that the adsorption rate constant of the labeled phages was comparable to that of unlabeled phages (**Fig S3**).

### Phage trajectory duration distributions reflect T4 attachment

We chose phage T4 and *E. coli* bacteria as our initial model because the interactions between these microbes have been widely studied (29). Moreover, T4 has been successfully labeled with a lysine-specific dye in prior studies, which allowed modification of an existing protocol (30). Importantly, more than one receptor is involved in T4 attachment to *E. coli* K-12, which permitted exploration of multiple bacterial mutants that should differ in terms of T4-attachment dynamics (31, 32).

In the MPA assay, a large field of view allowed us to obtain thousands of trajectories within minutes. To quantify the heterogeneity in the phage behavior (short, intermediate, and long trajectories in **Fig 1**D), we calculated the duration of each trajectory, and depicted the distributions of these durations. We note that trajectory duration (time that the phage remains within the microscopic focal view) is representative of the phage dwell time (time that the phage interacts with a bacterial cell). Initially, we focused on the trajectory duration distributions of phage T4 when interacting with two *E. coli* K-12 strains: wildtype bacteria (WT^T4^, WT cells carrying a kanamycin cassette in a pseudogene-see Materials and Methods for details) and a strain lacking outer membrane porin C (strain Δ*ompC*), the cognate receptor of phage T4. T4 is unable to form plaques on Δ*ompC* cells, and previous studies have indicated negligible adsorption rate for the phage on this host using the classical adsorption assay (32–34). Hence, we expected that individual phages would spend relatively less time interacting with Δ*ompC* cells, which should be reflected in the trajectory duration distributions. We observed that the numbers of trajectories with longer durations were considerably higher for phages interacting with WT^T4^ cells as compared to those interacting with Δ*ompC* cells (**Fig 2**A). The differences in these plots affirmed that quantitative differences in phage-bacteria interactions could be observed in our measurements at the single-phage resolution.

**Fig 2.**
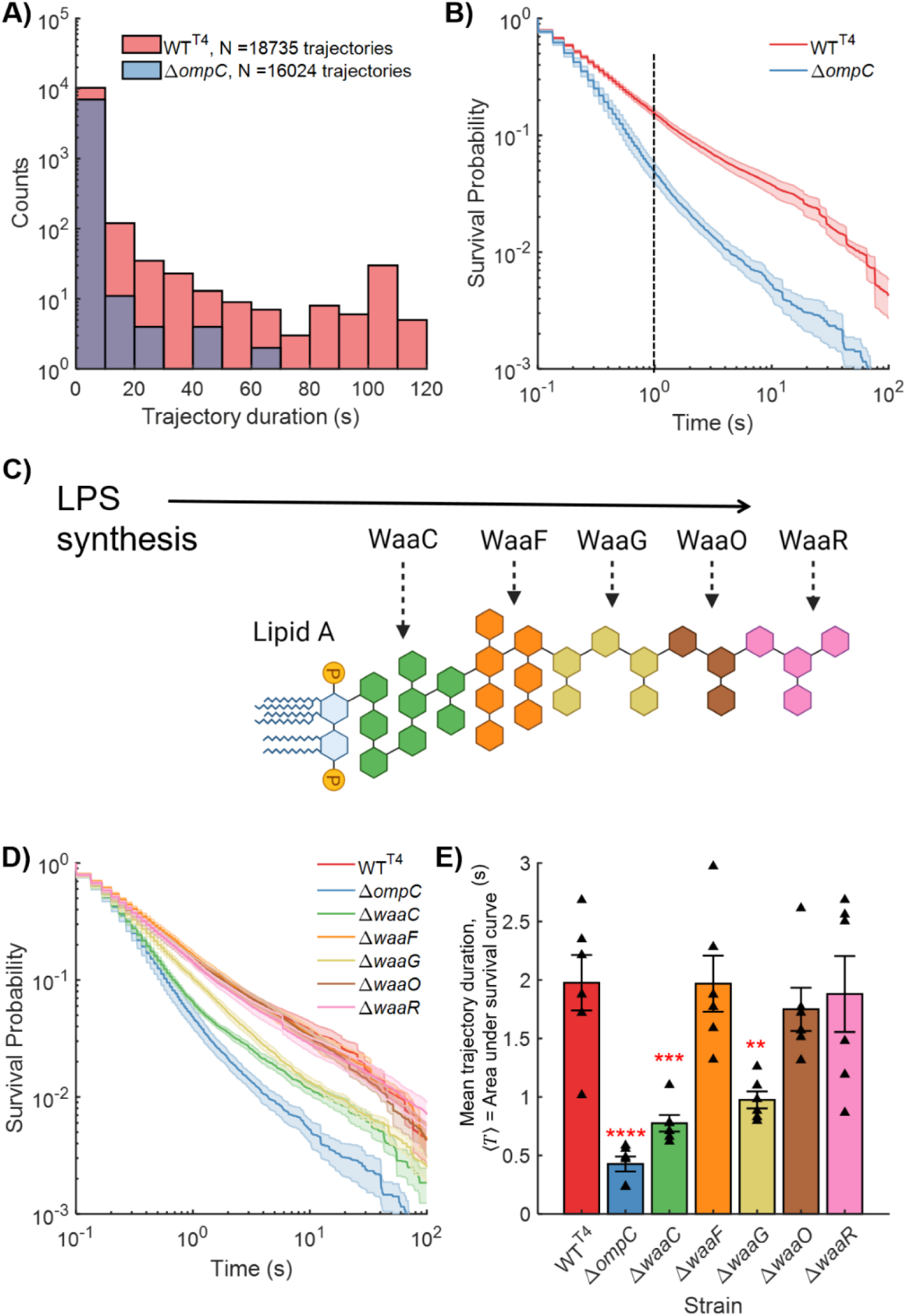
Phage attachment mutants yield distinct trajectory duration (dwell time) distributions. **(A)** Histograms are indicated for the trajectory durations of T4 phages interacting with wildtype cells (WT^T4^) and cells lacking the cognate T4 receptor outer membrane porin C (OmpC). **(B)** Survival probability *P*(*T* > *t*), the probability that a T4 trajectory is longer than *t*, was calculated for both strains indicated in **A**. At a representative *t* = 1 *s* (dashed black line), the probability that a T4 trajectory is longer than 1 s, *P*(*T* > 1) = 0.16 and 0.05 for WT^T4^ and Δ*ompC* cells respectively. **(C)** Lipopolysaccharide (LPS) structure has lipid A at its base. WaaC, WaaF, Waag, WaaO, and WaaR are enzymes that facilitate sequential addition of sugar layers to the LPS. Thus, a Δ*waaG* mutant has a shallow LPS, composed of only the layers indicated in green and orange colors. **(D)** Survival probability distributions for T4 interaction with each of the test strains are shown on the same plot. Total number of trajectories for each of the strains were: *N*_*WT*_*T*4 = 18735, *N*_Δ*ompC*_ = 16024, *N*_Δwaa*C*_ = 17416, *N*_Δwaa*F*_ = 20074, *N*_Δwaa*G*_ = 15717, *N*_Δwaa*O*_ = 16722, *N*_Δwaa*R*_ = 19762. **(E)** Mean T4 trajectory duration ⟨*T*⟩, calculated as the area under the survival probability distribution curves, is shown for each strain. Data indicate mean ± standard error for three biological replicates with two technical replicates each. Triangles correspond to ⟨*T*⟩ calculated for each experimental replicate. Statistical significance was assessed through ANOVA (P = 7×10^-7^) and post hoc Dunnett’s multiple comparisons test to compare each strain with WT^T4^. **P < 0.01, ***P ≤ 0.001, ****P < 0.0001.

Next, we calculated survival probabilities which allowed us to depict and compare the trajectory distributions for different cases with varying numbers of trajectories. We note that “survival” does not refer to the existence of either bacteria or phages; rather, “survival probability” is a quantity in statistics, defined as follows. Survival probability, *P*(*T* > *t*), represents the probability that a phage from the observed population exhibits a trajectory duration *T* that is longer than *t*. Thus, the “survival” in survival probability refers to the physical presence of a phage particle within the microscopic focal view. For instance, *P*(*T* > 0) = 1, because all observed trajectories have a non-zero duration. For large enough times *t* we observed that the survival probability of T4 with the two strains differs significantly, *P*(*T* > *t*)_*WT*_ ≫ *P*(*T* > *t*)_Δ*ompC*_ (**Fig 2**B, see dashed line). In other words, phage T4 exhibits a significantly higher probability of remaining longer in the microscopic field of view when interacting with WT^T4^ cells, as compared to interactions with Δ*ompC* cells. These results suggested that our measurements could distinguish between T4 phages interacting with adsorbable host cells versus hosts lacking the receptor required for phage binding.

Along with its primary receptor OmpC, T4 uses lipopolysaccharide (LPS) as a co-receptor (31, 32) for *E. coli* K-12. Hence, we examined five LPS synthesis mutants which were expected to produce different T4 attachment dynamics. We used mutants Δ*waaC*, Δ*waaF*, Δ*waaG*, Δ*waaO*, and Δ*waaR*, which lack the genes encoding the enzymes that facilitate the addition of various sugar molecules on the base lipid A (**Fig 2**C). Due to varying outermost sugar moieties, these mutants feature LPS layers that differ in total depth. While phage T4 forms plaques on each of these strains which have OmpC present, the attachment dynamics vary owing to these differences in LPS composition (32). We performed MPA assays with T4 and each of the LPS mutants. The survival probability distributions for T4 on all test strains are shown in **Fig 2**D. Survival probability distributions for individual replicate experiments are included in the Supplementary Text (**Fig S4**). We expected that the survival probability of T4 trajectories would be reduced when interacting with mutants featuring shallower LPS, as compared to WT^T4^ cells, but that all these survival probabilities should exceed that measured for Δ*ompC*, the un-adsorbable strain (32). The distributions for the shallowest (Δ*waaC*) and third-shallowest (Δ*waaG)* LPS-synthesis mutants were indeed intermediate to those for WT^T4^ and Δ*ompC*. Curiously, the survival probability for the second-shallowest LPS-synthesis mutant, Δ*waaF*, was higher than expected (32), comparable to that for WT^T4^. To investigate this, we sequenced these Keio-collection strains. Analysis of whole genome sequencing data revealed that the Δ*waaF* strain does not feature a true *waaF* deletion (see **Supplementary Note 1**). From this analysis, we concluded that the strains except Δ*waaF* can be reasonably considered as true deletions.

Finally, we calculated average trajectory duration (area under the survival probability distribution; **Supplementary Note 2**), which corresponds to the average time that a phage spends within the microscopic focal field of view, likely interacting with bacterial cells. Thus, the average trajectory duration represents a readout of the average dwell time. We observed that the average trajectory duration was shortest for Δ*ompC* cells, intermediate for the LPS synthesis mutants with the shallowest LPS (Δ*waaC*, Δ*waaG*), and longer for WT^T4^ cells as well as other LPS synthesis mutants (Δ*waaO*, Δ*waaR*; **Fig 2**E).

### MPA (microscopic) assay outcomes correlate with classical adsorption assay outcomes

Next, we performed classical adsorption assays to compare with the microscopy results. We hypothesized that the mean trajectory duration extracted from the MPA assay (presumably indicating more irreversible adsorption) should correlate with the adsorption rate constant in the classical adsorption assay (**Fig 3**A, also suggesting more irreversible adsorption).

**Fig 3.**
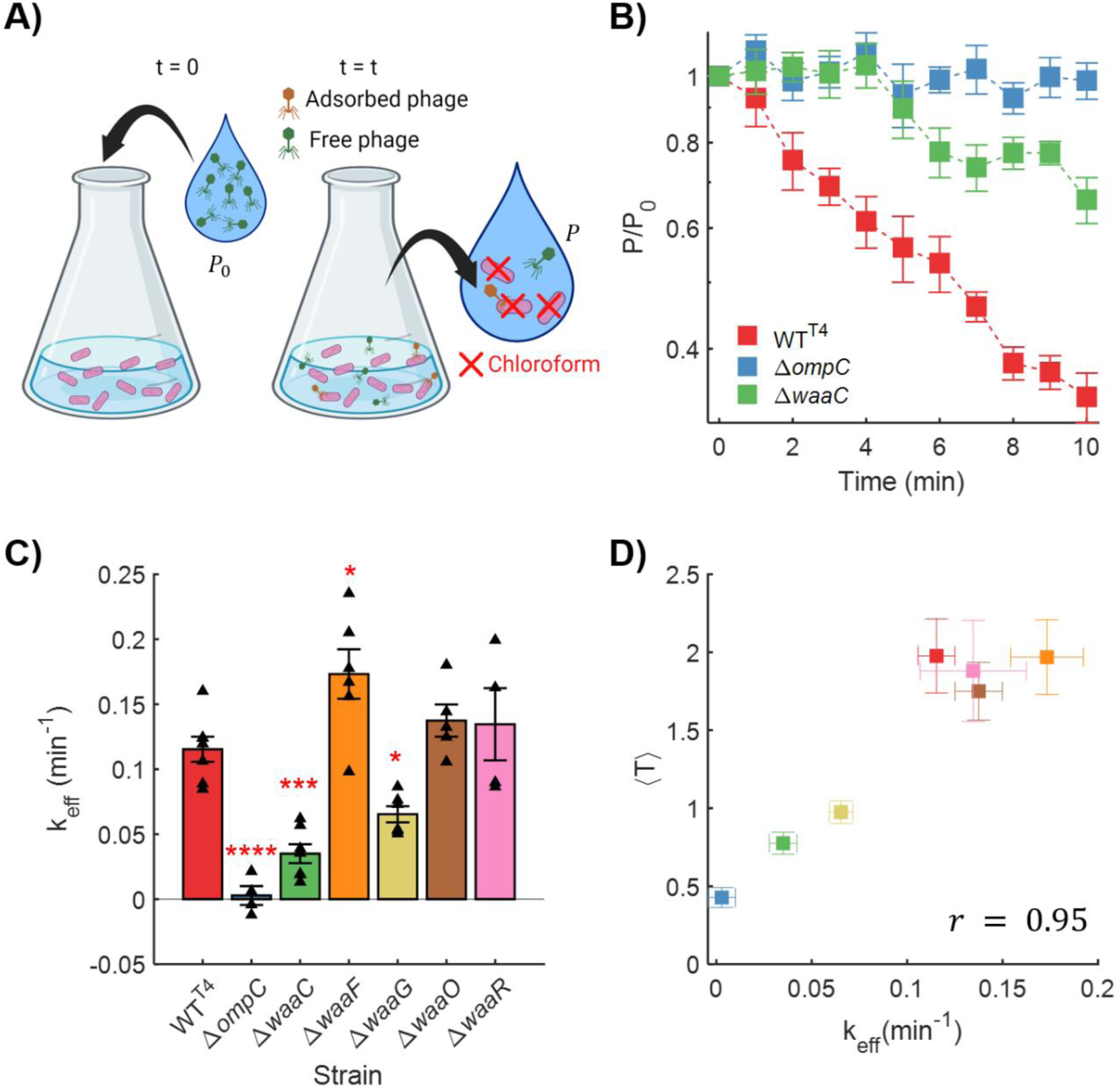
Trajectory duration distributions correlate with classical adsorption rate constant. **(A)** A schematic describing the classical adsorption assay: a known number of phages are added to an exponentially-growing bacterial culture at time ***t*** = **0**. Aliquots are obtained from the mixed culture at different time-points (e.g., every 1-2 min), followed by isolation and enumeration of free (unattached) phages. **(B)** Example classic adsorption traces are shown as fraction of free phages over time: the number of unattached T4 phages depletes fastest when interacting with WT^T4^ cells, slower in case of Δ*waaC* cells, and no apparent depletion is observed for Δ*ompC* cells. Mean ± SE at each time-point is indicated for each strain. A least-squares fit (Equation 1) to the classic adsorption curve is performed to calculate the adsorption rate constant. **(C)** Mean ± SE of effective adsorption rate constant ***k***_***eff***_, calculated from individual experimental replicates (triangles) is plotted for each strain. Statistical significance was assessed through ANOVA (P = 1.1×10^-9^) and post hoc Dunnett’s multiple comparisons test to compare each strain with WT^T4^: *P < 0.05, ***P ≤ 0.001, ****P < 0.0001. **(D)** The outcome of the microscopy assay, average trajectory duration ⟨*T*⟩ (**Fig 2E**), is plotted against the effective adsorption rate constant obtained via the classic assay (**Fig 3C**). Pearson correlation test revealed a high amount of linear relationship in these data with correlation coefficient *r* = 0.95 and P-value = 7.5×10^-4^ (calculated using a Student’s *t* distribution).

The fraction of free phages over time, i.e., the classic adsorption curves for T4 attachment to cells of WT^T4^, Δ*waaC*, and Δ*ompC* strains are shown in **Fig 3**B. The results corroborated the trajectory duration distributions : the depletion of free phages due to cell attachment is fastest for WT^T4^, intermediate for Δ*waaC*, and slowest (negligible) for Δ*ompC* (**Fig 2**D), consistent with the survival probability of T4 being lowest for Δ*ompC*, intermediate for Δ*waaC*, and highest for WT^T4^ (**Fig 3**B). Classic adsorption curves from individual biological replicates are included in the Supplementary Text (**Fig S5**).

We calculated the effective adsorption rate constant for each strain from each classic adsorption curve by assuming a pseudo-first order association between bacteria and phages (22, 35), and obtaining least-square fits for effective adsorption rate constant *k*_*eff*_:

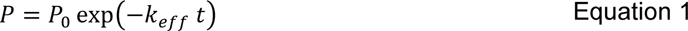

where the number of PFUs drops from initially *P*_0_ to *P* at time *t*. The assumptions and implications underlying this adsorption model are discussed in **Supplementary Note 3**. The resultant effective adsorption rate constant values for each strain are indicated in **Fig 3**C. As expected, the mean adsorption rate constants for Δ*ompC*, Δ*waaC*, and Δ*waaG* strains were significantly lower than that for WT^T4^ cells.

Next, we examined to what extent the average trajectory duration for each strain in the MPA assay (**Fig** 2D) correlated with the adsorption rate constant in the classic assay (**Fig 3**C). Plotting one against the other revealed a monotonic relationship with a correlation coefficient of 0.95 (**Fig 3**D) (P = 7.5×10^-4^; Pearson correlation test, see Materials and Methods). Thus, the average trajectory durations strongly correlated with the classically determined adsorption rate constant.

We concluded that the MPA assay outcomes strongly corroborated those obtained using classical adsorption methods, with significantly reduced labor and the added benefit of measuring heterogeneity at the single-phage resolution as opposed to an ensemble estimate.

### MPA assay is generalizable to other phages and bacterial species

We hypothesized that our phage-labeling protocol and time-lapse imaging technique were versatile; i.e., we could use the MPA assay to quantify attachment of various phages to bacterial cells of differing species. To test this hypothesis, we performed phage-labeling and MPA assay with a diverse set of phages that were specific to *E. coli* and to other host bacteria, and which varied in morphology and size (i.e., capsid diameter and tail length) (**Table 1**). Phage size is an important factor that determines the number of dye molecules attached to each phage particle and hence the intensity of the fluorescence signal emitted by a phage particle. We therefore were able to visualize all phages but the smallest one (coliphage ϕX174) using our widefield fluorescence microscopy setup.

**Table 1.** Morphology and microscopic outcomes for the phages used in this study.

| Phage | Host | Capsid Diameter (nm) | Particle Length (nm) | Fluorescent foci visibility |
| --- | --- | --- | --- | --- |
| T4 | *E. coli* | 120 | 80 | ✓ |
| $\lambda$ | *E. coli* | 60 | 150 | ✓ |
| T5 | *E. coli* | 90 | 160 | ✓ |
| M13 | *E. coli* | 7 | 880 | ✓ |
| T2 | *E. coli* | 114 | 115 | ✓ |
| T7 | *E. coli* | 60 | 20 | ✓ |
| $\phi X174$ | *E. coli* | 25 | 25 | ✗ |
| P1 | *E. coli* | 85 | 228 | ✓ |
| A1-1 | *S. flexneri* | 97 | 143 | ✓ |
| OMKO1 | *P. aeruginosa* | 139 | 210 | ✓ |

We conducted the MPA assay with phages listed in Table 1 utilizing cognate (adsorbable) hosts as well as strains or conditions known to disrupt phage attachment. As test strains, especially while experimenting with coliphages, we made extensive use of the Keio collection of single-deletion *E. coli* K-12 mutants and especially its parent strain BW25113, referred to as WT (wildtype) (36). The resultant trajectory duration distributions for all the experimental phages are indicated in **Fig 4**. Data for average trajectory duration 〈*T*〉, the key parameter reflecting phage-affinity to bacteria, are shown as inset results for each experimental case.

**Fig 4.**
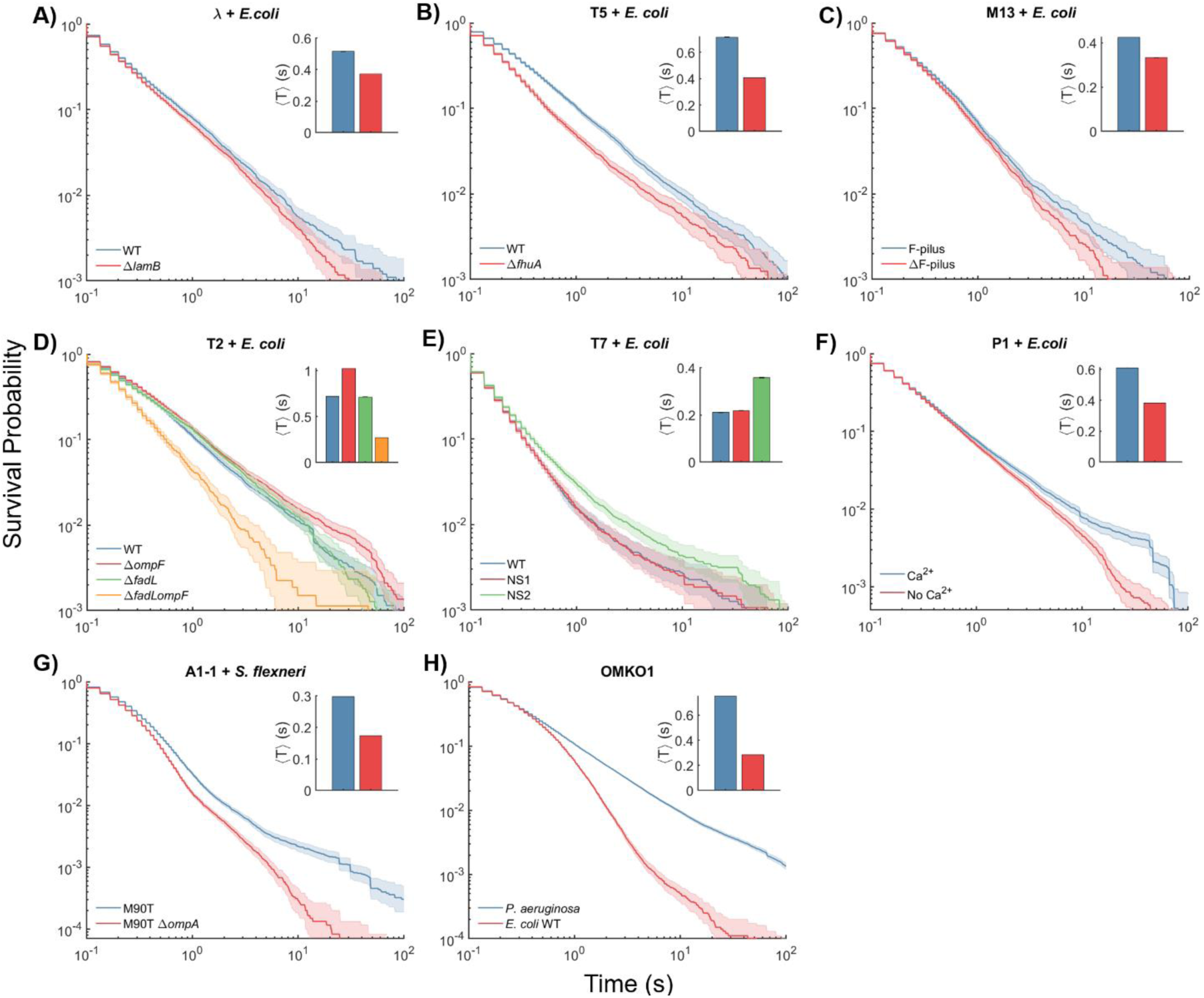
MPA assay can be generalized for other phage and bacterial species. Survival probability distributions for the trajectory durations are plotted for each phage and strain of its host bacteria. The average trajectory duration 〈*T*〉 for each strain is shown as inset, with the same color scheme as the survival probability distributions and error bars calculated through bootstrapping. **A-F** feature coliphages (phages that infect *E. coli*), interacting with bacterial cells in known conditions favorable or unfavorable for phage attachment. **G-H** feature visualization of phages that infect two biomedically relevant species, *S. flexneri* and *P. aeruginosa*.

Phage λ is an extensively studied coliphage model (37). It uses the maltoporin LamB as a receptor (37). We tracked single λ phages interacting with WT cells as well as a mutant lacking *lamB.* Phage T5 uses the outer membrane protein FhuA as the receptor (34, 38). We visualized attachment of T5 phages to WT and an *fhuA* knockout. Filamentous phage M13 infects *E. coli* cells through the F-pilus (39). We carried out MPA assays with strain CSH22 which carries the pilus as the “F-pilus” strain and WT as the strain lacking the F-pilus or “ΔF-pilus” strain. In each of these cases, results confirmed that the trajectory survival probability distributions were skewed towards lower values and the average trajectory duration was considerably lower for the receptor-less mutant as compared to WT (**Fig 4**A,B,C).

Coliphage T2 is morphologically similar to T4; however, it can use either of two receptors, OmpF and FadL, to infect host cells (34, 40, 41). We used WT and mutants lacking either or both of the genes encoding OmpF and FadL as our test strains for T2-MPA assay. We observed that the trajectory duration distribution is skewed towards lower values for the double mutant (**Fig 4**D), indicating considerable differences in the adsorption to this mutant as compared to the other strains, in agreement with classical adsorption assay measurements reported earlier (34).

Phage T7 binds to LPS of *E. coli* host cells, and cannot form plaques on Novobiocin-supersensitive (NS) *E. coli* mutants NS1 and NS2 (42). For T7-MPA assay, we used WT, NS1, and NS2 as test strains. The mean trajectory duration was higher for the NS2 strain as compared to WT and NS1 (**Fig 4**E). These data suggest the explanation that T7’s infectivity of the NS strains is likely not hampered at the attachment stage of the infection process (in fact, attachment to NS2 appears to be stronger than to WT). Instead, the later stages of infection (e.g., genome injection, replication) may be responsible for the T7-resistance of NS strains, which we are unaware has been explored to date.

Phage P1, a phage commonly used as an *E. coli* genome manipulation tool, requires the calcium cation (Ca^2+^) for successful adsorption (43). We carried out the MPA assay with labeled P1 attaching to WT cells with or without the addition of 5 mM Ca^2+^. Data confirmed the trajectory duration was lower in the absence of Ca^2+^ (**Fig 4**F), indicating reduced adsorption.

Next, we evaluated our method with phages infecting two key bacterial species of biomedical significance: *S. flexneri* and *P. aeruginosa* (**Table 1**). Phage A1-1 infects *S. flexneri* cells through outer membrane porin A (OmpA) and LPS as co-receptors(44). We successfully labeled A1-1 and performed the MPA assay with host strain *S. flexneri* M90T (WT) and its *ompA*-knockout mutant. As expected, the trajectory durations were lower in case of Δ*ompA* (**Fig 4**G). For *Pseudomonas*-infecting jumbo-phage OMKO1, which has been successfully used in therapy (45, 46), we used *P. aeruginosa* PAO1 strain as the adsorbable host and *E. coli* WT as negative control for attachment. Results confirmed that the phage trajectory durations were significantly lower when interacting with *E. coli* cells (**Fig 4**H).

## Discussion

We present a novel assay to quantify attachment of phages to bacteria by tracking individual particles using widefield fluorescence microscopy, which we term as the Microscopic Phage Adsorption (MPA) assay. Previous studies have used multiple approaches to perform dynamic visualization of individual phages, including genetic fusion of fluorescent proteins to capsid constituents (24, 47, 48), streptavidin-biotin conjugation to functionalize dye molecules to biotinylated sites on phage particles (49), nanoparticles (49), or quantum dots to combine with phage capsids (50), as well as dyes that conjugate with genetic material (51, 52) or phage surfaces (30). Circumventing the challenges associated with phage-genome engineering, we sought a method applicable across various phage species. Therefore, we opted for a dye utilizing TFP (tetrafluorophenyl) ester chemistry – an improvement over the popular NHS (*N*-Hydroxysuccinimide) ester chemistry – capable of specifically targeting all lysine residues exposed on any biological surface. Seeking the simplest and least time-consuming experimental method, we developed a minimal protocol without extra cleaning steps such as PEG-precipitation, CsCl ultracentrifugation, or gel-based purification (47, 48). Our method yielded viable phages that were capable of lysing cells. Except for a key limitation related to phage size (**Table 1**) discussed below, our labeling and single-particle-tracking approach worked for various phages with different morphologies (**Fig 4**). Notably, phage M13 has a distinct filamentous shape, differing significantly from the head-tail morphology of many other known phages. Despite this difference, our approach enabled visualization and quantification of M13 phages adsorbing to hosts. Thus, our visualization technique is applicable to diverse phage morphologies.

Importantly, our technique provides an alternative to the laborious and low-throughput classical adsorption assays. These plate-based assays involve preparation of various types of consumable reagents and culture media used in different forms, especially 1.5% agar plates, 0.75% top agar, and liquid media housed in sterile glassware. Several manual experimental steps are also involved, such as determining concentration of the phage stock and preparing the desired initial (t = 0) phage concentration; pre-warming plates and top agar; preparing chloroform or filters for separation of free phages from bacteria in each aliquot; pouring the aliquoted phages mixed with fresh bacteria and top agar on agar plates; and finally counting the PFUs on each plate (20, 53). Also, 12-48 hours of incubation time (depending on the bacterial species) is required for visible plaques to form on the plates. Due to these many steps and abundant consumables, researchers often must limit the number of aliquoted time-points to achieve practicality (54, 55), which can reduce empirical accuracy. After optimizing these experiments, an ensemble estimate of the adsorption rate constant is calculated based on a model (20, 22, 53). Our approach instead allows direct model-free observation of phage dwell time (time spent by a single phage interacting with bacteria) for each individual phage particle, while avoiding the laborious steps and incubation-wait-time described above. We utilized the simplest of the fluorescence microscopy techniques, and the results strongly correlate with the traditional measurements of adsorption rate constants (**Fig 3**). Therefore, we anticipate that the MPA assay—which provides a more robust measure of phage-attachment at single-virion level— could become widely useful for researchers in phage biology who have access to a fluorescence microscope.

Phage interaction with bacterial cell surfaces can include reversible and irreversible attachment steps (18, 22, 56). Models in the literature have inferred the rates associated with reversible and irreversible adsorption from classical assay outcomes (22). However, the actual fraction of phages that attach to bacterial cells (also termed adsorption efficiency (57)) as well as numbers of successful versus unsuccessful collisions with host cells is impossible to estimate without dynamic probing of single phages. While faster and more efficient adsorption could provide proximate benefits to the phage, maximizing attachment may not always be an ideal phage ecological strategy, as theorized by Gallet et al (58). In a hypothetical scenario where each virion from a homogeneous (single genotype) phage population can attach to a bacterial cell upon collision, the phage population would deplete rapidly. However, if relatively fewer of these isogenic particles attached to cells, a fraction of the phage population could sample other host types in the environment to possibly expand niche space (e.g., cells capable of supporting equal or greater phage production compared to the typical host). A similar argument can be made about temporary phage dormancy: if phages can temporarily become attachment-deficient, they can improve their chances of sampling other host types (59). Thus, reversible binding or phage dormancy afford the possibility for phages to explore other host niches, promoting greater overall growth of the phage population containing a single virus genotype. Our observations showed that the vast majority of phage encounter with cells do not result in permanent attachment, an outcome that cannot be observed directly using the classic approach. Instead, our results revealed that a majority of trajectories exhibit short or intermediate durations (**Fig 2**, **Fig 4**). These results provide considerable support for Gallet et al.’s theory (58), demonstrating that a vast majority of contact events between phages and cells do not lead to permanent attachment, allowing the phages to disperse and propagate efficiently if suitable alternative host cells are encountered. We note that this mechanism does not necessarily indicate an adaptive trait. That is, reduced variation among particles that minimizes reversible binding may be a property that cannot be improved through natural selection. This constraint could however provide an occasional useful outcome, if a phage variant reversibly binds to the typical host and happens to infect a new or altered host-cell type encountered in the environment. We believe that our approach is better suited to exploring such questions of efficiencies in reversible and irreversible binding in phages (whether molded through selection versus governed by stochastic processes), relative to the traditional method.

A prime application of the microscopy-based MPA assay is the evaluation of different candidate phages considered for use in applications such as phage therapy. Adsorption rate could be a key trait when determining the preferred phage candidate in certain applications, because faster (slower) attachment should allow a larger (fewer) number of target bacterial cells to be infected per unit time, all else being equal (12). For example, the best phage or combination (cocktail) of phages needs to be evaluated before designing and deploying phage treatment delivered to the patient. In a personalized medicine approach of phage therapy, the MPA assay can be conducted within minutes to screen a pre-labeled library of phages for attachment to bacteria isolated from the patient. By comparing the trajectory duration distributions for different phage candidates on the same target strain of host bacteria, a decision concerning the phage(s) that are most efficient at attaching to host cells can be made.

A key limitation to our approach is represented by the size of the viral particle. We successfully labeled and observed phages with characteristic sizes ranging from 25 to 139 nm capsid diameter, and those with 25 to 880 nm particle length (**Fig 4**), which span the size dimensions that appear typical for known phages. Phages spanning at least ∼50 nm in one dimension are expected to have hundreds of exposed lysine residues, allowing for the formation of a bright image on the microscope due to the binding of numerous dye molecules. In contrast, some phages have smaller capsid diameter as well as shorter particle length, resulting in a lower surface area and hence possessing relatively fewer lysine residues exposed on a single virion. For instance, we attempted to label ϕX174, a phage with a capsid diameter of 25 nm, and length of 25 nm (60). However, the fluorescence signal emitted by the labeled phages was insufficient to visualize them accurately, even at the maximum intensity of our LED light source. We believe this challenge can be addressed by using a stronger illumination source such as a LASER, and this idea warrants further evaluation in future work. Albeit, when considering applications such as phage therapy, this discussion may be of little concern, as the typical phages used in therapy are ∼100 nm or larger in size (46, 61).

One of our major concerns was potential blockage of crucial lysine residues in the phage tail fiber by the dye molecules, which could impact the interaction with the host surface (62). Our control experiments revealed that host cells were successfully killed by the labeled phages (Movie S1, **Fig S1**). Additional experiments also suggested that there is no significant difference between the ensemble adsorption rate constant of labeled and unlabeled phage T4 particles (**Fig S3**). Finally, our investigations involving various known bacterial mutants yielded noticeable differences in attachment that aligned with the classical adsorption assays performed with unlabeled phages. Thus, we conclude that the lysine-specific labeling approach is suitable for conducting comparative experiments on phages and bacteria, especially for high-throughput characterization of adsorption for phage and bacteria libraries. However, we recognize that our approach might not capture finer details in phage-adsorption dynamics that involve certain lysine residues. To design experimental studies of precise biophysics of phage attachment, this limitation can be addressed by labeling the genome of phages with DNA-specific dyes or opting for fluorescent-protein fusions. We anticipate that such studies would require a spatial resolution which is not afforded by our deliberately chosen simplest fluorescence microscopy technique.

Our single-phage visualization method opens up new avenues to investigate the dynamic steps involved in phage interactions with the host-cell surface. For example, common biophysical parameters quantifying the target search of microscopic particles include diffusion coefficients as well as the relative occupancies of different diffusive states and transition rates between these states (63–66). With our microscopy approach, phage interactions with cells can be recorded at high spatiotemporal resolutions and biophysical parameters of the phage target search can be determined using various analyses reported in the literature (63, 65, 66). The generalizability of our phage labeling approach will allow the study of differences in target search dynamics as consequences of diverse viral morphology, phage species, receptors, and bacterial species. Moreover, our approach could be used to explore how phenotypic variation among particles in a phage population changes over evolutionary time as viruses evolve differences in using cell receptor(s), such as longitudinal analyses of samples from experimental evolution studies that test ideas analogous to emergence of viruses on novel host types. Beyond phages, our approach combining biomolecule labeling, microscopy, particle tracking, and trajectory duration analysis can potentially be employed to compare the dynamics of other viruses (e.g., mammalian or plant viruses) or drug molecules binding to their respective targets. This broader application widens the scope of our approach, making it a versatile tool for investigating a wide range of biomolecular interactions.

## Materials and Methods

### Bacteria and phage strains

All *E. coli* K-12 bacteria and phages of *E. coli* used in this study (**Table 1**, **Table 2**) were kindly provided by J. Wertz at the Coli Genetic Stock Center (CGSC) at Yale University, which is now *E. coli* Genetic Resource Center (https://ecgrc.net/). Strain BW25113 was used as the wildtype (WT) *E. coli* host for phages T2, T5, T7, P1 and λ, whereas derivatives of BW25113 in the Keio collection were mutants that contained individual gene knockouts and were used as challenge hosts due to genetic changes in cell-surface receptors (**Table 2**). For phage T4, a mutant with knockout in pseudogene *icdC* was used as a proxy for wildtype (WT^T4^). For phages M13, A1-1, and OMKO1, strains *E. coli* CSH22, *S. flexneri* M90T, and *P. aeruginosa* PAO1, respectively, were used as wildtype hosts; either a knockout of these hosts or strain BW25113 (WT) was used as the challenge host lacking the cellular receptor for each of these viruses (**Table 2**). Phages A1-1 and OMKO1 have been described previously (44, 67).

**Table 2.** Bacterial strains used in this study. (*Eschericia coli K-12* unless otherwise specified). * CGSC# refers to the catalog number at the Coli Genetic Stock Center at Yale University, now available through *E. coli* Genetic Resource Center (https://ecgrc.net/).

| Phage | Strain abbreviation | Strain number* |
| --- | --- | --- |
| T4 | WT<sup>T4</sup> | Keio collection $\Delta icdC$, CSGC #11220 |
| | $\Delta ompC$ | Keio collection, CSGC #9781 |
| | $\Delta waaC$ | Keio collection, CSGC #11805 |
| | $\Delta waaF$ | Keio collection, CSGC #10648 |
| | $\Delta waaO$ | Keio collection, CSGC #10652 |
| | $\Delta waaR$ | Keio collection, CSGC #10651 |
| | $\Delta waaG$ | Keio collection, CSGC #10655 |
| T2 | WT | Keio parent strain BW25113 |
| | $\Delta ompF$ | Keio collection, CSGC #8925 |
| | $\Delta fadL$ | Keio collection, CSGC #9875 |
| | $\Delta fadLompF$ | JM1101, CGSC #5844 |
| T5 | WT | BW25113 |
| | $\Delta fhuA$ | Keio collection, CSGC #8416 |
| T7 | WT | BW25113 |
|  | NS1 | CGSC #6517 |
|  | NS2 | CGSC #6518 |
| M13 | F-pilus | CSH22 |
| | $\Delta F$-pilus | BW25113 |
| P1 | Ca<sup>2+</sup> | BW25113 |
|  | No Ca<sup>2+</sup> | BW25113 |
| $\lambda$ | WT | BW25113 |
| | $\Delta lamB$ | Keio collection, CSGC #10877 |
| A1-1 | M90T | *Shigella flexneri* M90T |
| | M90T $\Delta ompA$ | *Shigella flexneri* M90T $\Delta ompA$ |
| OMKO1 | *P. aeruginosa* | *Pseudomonas aeruginosa* PAO1 |
|  | *E. coli* WT | BW25113 |

Lysogeny broth (LB; 10 g tryptone, 5 g yeast extract, 10 g NaCl per L) was used for liquid cultures, 1.5% agar plates, and 0.75% top agar for the double-layer method of phage propagation. With exception of experiments using phage T4 and phage P1 interacting with WT in absence of Ca^2+^, LB was supplemented with 25 μg/mL thymine, 10 mM MgSO_4_, and 5 mM CaCl_2_. 0.1% D-(+)-Maltose was further supplemented to the growth and experimental media for phage λ. Overnight cultures were initiated from colonies grown on solid (1.5%) LB agar and placed into LB liquid medium, incubated with shaking (175–200 RPM) at 37°C. Serial culture was conducted by diluting 0.1 mL overnight culture into 10 mL fresh LB medium. Densities of bacteria were determined by plating dilutions from cultures onto agar plates, to estimate colony-forming units (CFU) per mL. Bacterial stocks were stored in 25% glycerol at -80°C. Phages were enumerated (plaque-forming units [PFU] per mL) using the standard double-layer agar method where viruses form visible plaques on confluent lawns of wildtype host bacteria within 0.75% top agar, overlayed on 1.5% agar in the bottom layer. High-titer stocks (lysates) of phages were grown using the double-layer agar method: top agar layers with a lacey plaque-pattern were suspended in PBS, centrifuged, and filtered (0.22 μm) to remove bacteria and obtain a cell-free lysate.

### Phage labeling with fluorescent dye

Fluorescent labeling of phage T4 with Alexa Fluor^®^ 488 NHS Ester dye was previously described (30). An improved dye-amine conjugate, TFP (tetrafluorophenyl) ester, a more hydrolysis-resistant amine-reactive ester as compared to NHS (succinimidyl) ester, was utilized in the current study: AZDye™ 488 TFP Ester from Fluroprobes (Catalog No. FP-1026, Alexa Fluor^®^ 488 TFP Ester equivalent) was used in all labeling protocols. While dye wavelength (488 nm) was chosen for compatibility with our in-house fluorescence microscopy filter sets, we have successfully used the same labeling protocol for dyes excitable at other wavelengths or offering conjugation-chemistry other than amine-reactive esters. Amicon^®^ Ultra-4 centrifugation filter columns with a cutoff size 100 kDa were used to transfer ∼10^8^-10^9^ phages from a lysate to phosphate buffer saline (PBS, pH 7.4, Gibco^TM^). The 100 kDa cutoff was sufficient to filter out components in the media while retaining phages. Transferring phages to PBS ensured robust dye-conjugation as competing growth-media components carrying lysine molecules were removed before introducing the dye. The phages were then conjugated with 0.5 mg/mL dye via shaking (500 rpm) incubation at room temperature for 3 hours, followed by static incubation at 4 °C overnight. Excess unbound dye was removed via centrifugation (4-5 washes) using another Amicon^®^ Ultra-4 filter. Labeled phages were stored at 4 °C until imaging. Phages were imaged within three weeks of conjugation in order to avoid lower fluorescence signal due to dye degradation.

### Microscopy (MPA assay)

Single phages were visualized via epifluorescence microscopy at high spatiotemporal resolution. A Nikon Ti-E inverted microscope equipped with a 1.40 NA objective with 100× magnification, perfect focusing system, and temperature-controlled chamber was used. The temperature was maintained at 34 °C throughout the experiments. Tunnel slides (glass slides with coverslips affixed at edges via double-sided adhesive tape) were used to prepare samples. In the experiments measuring free phage diffusion, the objective was focused ∼10 μm above the coverslip surface. For live phage-bacteria interaction assays (the MPA assay), 0.1% poly-L-lysine was first introduced to the tunnel slide, which was then incubated upside down for at least 5 and up to 30 minutes, to coat the coverslip. Exponential-phase bacteria from freshly-grown cultures were washed twice in PBS, concentrated, and flowed into the tunnel slide, followed by a 10-minute upside down incubation to adhere and immobilize the bacteria. After washing away unadhered bacteria with the growth medium, fluorescently-labeled phages suspended in LB growth medium (as it contains attachment cofactors such as tryptophan (17, 68)) with the appropriate supplements described above, were introduced to the tunnel slide. The tunnel-openings were immediately sealed with VALAP (equal parts Vaseline, Lanolin, and Paraffin Wax) to avoid evaporation and flows, and the sample was mounted onto the microscope for imaging. The objective was focused ∼0.5 μm above the coverslip surface, identified via visualization of phages immobilized at (attached to) the poly-L-lysine-coated surface. A snapshot of underlying bacterial lawn was recorded in the phase-contrast channel. Finally, videos of phages attaching to bacteria were recorded in the fluorescence channel at a high temporal resolution of 30 frames per second. To ensure comparability between experiments, experimental replicates of the same phage interacting with different host/non-host strains were obtained on the same day.

### Analysis of microscopy videos to obtain single phage trajectories

From videos of fluorescently labeled phages recorded at 30 frames/s, trajectories of single phage particles interacting with host cells were obtained using custom-written MATLAB algorithms for particle tracking, similar to algorithms described earlier (69–71). Our algorithm is described below, which is equivalent to popular particle tracking programs such as `trackpy` in Python, and `TrackMate` in ImageJ/Fiji:

### Detection

Each image was smoothed with a Gaussian kernel with a standard deviation (std. dev.) significantly less than the pixel-size of a feature (bright image formed by a single phage). A background image was generated by smoothing the image with a std. dev. significantly greater than the feature. The second image was subtracted from the first image, resulting in significantly sharper features. As most of the pixels did not contain a phage, the histogram of individual pixel intensities (calculated by pooling several images) was fit in the vicinity of its peak with a Gaussian distribution to extract the mean and std. dev. of the noise pixel intensities. All pixel intensities that were less than 2 noise std. dev. above the noise mean were set to zero. Most of the patches of non-zero pixels left in each frame contained phages. Patches consisting of a single non-zero pixel were set to zero intensity, because phages typically occupy several pixels. Patches in which the highest-intensity pixel was less than 5 noise standard deviations above the noise mean were also set to zero, only leaving pixel patches that contained phages in focus. Finally, the phage-positions were refined to sub-pixel resolution by fitting a 2D Gaussian profile to each phage image (72).

### Tracking

As a first pass, starting from the first frame, for each detection in the frame, the closest detection in the next frame (frame gap 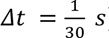) within a search radius of 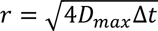 was found, where *D*_*max*_ was set to 10 μm^2^/s, which is significantly larger than 3 μm^2^/s, the typical mean diffusion coefficient of a free phage particle (**Fig 1**). Then, for each detection in frame *i*, the closest detection in frame *i* + 1 within search radius *r* was considered the same phage and linked into a trajectory. This process was repeated for each phage in the frame, and then for all frames, in chronological order. Trajectories that lasted only a single frame were considered false positive detections and were removed.

The assignment procedure described above could leave gaps in the trajectory of a given phage if it disappeared from the field of view for a few frames, for example by diffusing out of the depth of field and immediately diffusing back in. Linking trajectories across these gaps allowed a more accurate estimation of Dwell time. To close such gaps, trajectories were looped over from longest to shortest for a given gap size of *k* frames. For each trajectory, trajectories that started *k* frames after (before) the current one within a distance 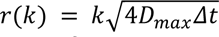 of the current trajectory’s end (start) were searched. If multiple nearby trajectories were found, the closest one was linked to the current trajectory. The loop was then continued to the next trajectory. Trajectories for which links were made were then revisited. Once no more links were made at gap size *k*, the gap size was increased by 1 frame. This process was repeated up to a maximum of 3 frames (0.1 s). Finally, trajectories that lasted only two frames were considered false positives and were removed.

### Multiplicity of infection (MOI) in microscopy assays

Multiplicity of infection (MOI) was calculated for each dataset (i.e., one snapshot of the immobilized bacterial cells + movie of fluorescently labeled phages interacting with those bacteria). Number of cells, *N*_*b*_, was calculated from the phase-contrast image of bacteria. Number of phages, *N*_*p*_ was calculated as median number of detections in the first hundred frames of the fluorescent-phage movie. MOI was calculated as

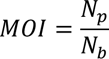

### Trajectory duration distributions and survival probabilities

Dwell time *T*_*i*_ for each trajectory *i* represented the duration of the trajectory. Histograms were plotted for the linear array *T* for each condition (WT and Δ*ompC*) as counts of elements in each bin. The bin size for the plot (**Fig 2**A) was chosen with a bin width of 10 s.

The survival probability for a given Dwell time (trajectory duration) array *T*, was calculated as

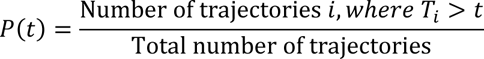

### Correlation between microscopy and classic assays

The correlation coefficient *r*, also known as Pearson correlation coefficient, was calculated as

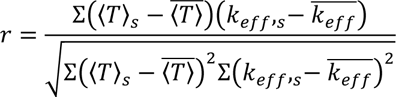

where, ⟨*T*⟩_*s*_ is the mean trajectory duration for strain *S* (mean value from all replicates, **Fig 2**D), *k*_*eff*_,_*s*_ is the mean effective adsorption rate constant for strain *S* (**Fig 3**C), and 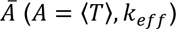 is the average value of individual ⟨*T*⟩_*s*_ and *k*_*eff*_, _*s*_ values. Strain *S* refers to each of the seven experimental *E. coli* strains discussed in **Fig 2** and **Fig 3**.

The corr algorithm in MATLAB was employed to calculate both the correlation coefficient and the corresponding p-value, which is computed using a Student’s *t* distribution for a transformation of the correlation.

### Classical adsorption assays

The adsorption assay used in this study was previously described (20). Briefly, cells were grown to OD_600_

∼0.5 and diluted to OD_600_ ∼0.1-0.2. A known concentration of phages (∼2-4×10^4^ PFU/mL) was mixed with bacterial cells taken from an exponentially-growing culture at *t* = 0. The mixture was grown with 34°C incubation, the same temperature as in microscopy experiments, with gentle shaking (60 rpm). Aliquots were sampled from the culture every minute for ten minutes. Each aliquot was immediately added to chilled tubes (4 °C) containing 3-4 drops of 100% chloroform and the tube was thoroughly vortexed. 4 mL of molten 0.75% top agar and 100 μL overnight bacterial culture were added to the tube, vortexed, and poured on 1.5% bottom agar plate incubated at 37°C. Plaques visible after 16-24 hours were counted to estimate the number of free (non-attached) phage particle concentration at each time point. The effective adsorption rate constant *k*_*eff*_ was calculated by obtaining least square fits for

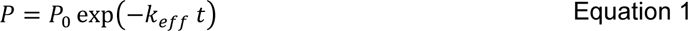

where *P*_0_ is the initial concentration (PFU/mL) of phages, and *t* represents time that it takes for the phage concentration to decrease from *P*_0_ to *P* (see **Supplementary Note 3** for derivation).

## Author Contributions

Conceptualization: JDA, TE, PET

Methodology: JDA

Investigation: JDA, TW

Visualization: JDA

Resources: TE, PET

Validation: JDA, TE, PET

Supervision: TE, PET

Writing—original draft: JDA

Writing—review & editing: JDA, TE, PET

## Acknowledgments

We thank Dallas Mould for careful and critical review of this manuscript. We thank John Wertz for providing the *E. coli* strains and coliphages used in this study as well as for commenting on the manuscript draft. Yale Quantitative Biology Institute members Henry Mattingly, Isabella Graf, Michael Blazanin, Asher Leeks, Albert Vill, Jeremy Moore, and Swayamshree Patra provided valuable discussions and feedback throughout the development of this work; we are grateful for their feedback. This study was funded by Yale University through Yale Center for Phage Biology and Therapy. TE acknowledges support from NIH (R01GM106189-09). JDA and PET acknowledge funding support from Howard Hughes Medical Institute Emerging Pathogens Initiative grant (30207345).

