## Supplementary Information for "Microscopic Phage Adsorption Assay: High-throughput quantification of virus particle attachment to host bacterial cells"

### Supplementary Note 1. Whole Genome Sequencing of Keio strains

| Strain | deleted genes* |
|---|---|
| WT$^{T4}$ | *icdC* |
| Δ*ompC* | *ompC* |
| Δ*waaC* | *[waaF],waaC,[waaL]* |
| Δ*waaF* | *[waaG],waaQ* |
| Δ*waaG* | *[waaG],[waaQ]* |
| Δ*waaO* | *waaR,[waaB]* |
| Δ*waaR* | *[waaY],waaJ,[waaR]* |

* Square brackets represent partial deletions and no brackets represent full deletions.

The sequence of LPS-related genes as in the MG1655 gemone on biocyc.org is shown below:

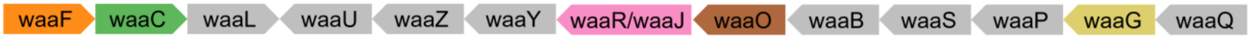


In the whole genome sequence of our ancestral strain BW25113 annotated with GenBank accession number CP009273, we observed the following sequence:

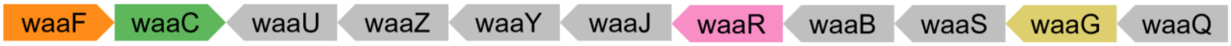


Mutations in each of our test strains were identified via the breseq mutant calling algorithm (1), and are listed in the table above. The differences in the waaJ-waaR-waaO region may explain the discrepancies in the final two rows of the above table: waaO and waaR are likely annotated as waaR and waaJ respectively.

### Supplementary Note 2. Area under survival curve represents mean trajectory duration

For individual trajectory duration $T$, survival probability as a function of time $t$:

$$F(t) = P(T > t)$$

then,

$$f(t) = -F'(t)$$

where, $f(t)$ is the probability density function, such that

$$\int_0^\infty f(t)\,dt = 1$$

Area under survival curve,

$$AUC = \int_0^\infty P(\tau > t)\,dt = \int_0^\infty F(t)dt$$

$$= \int_0^\infty F(t) \cdot 1\,dt$$

$$= t \cdot F(t) \Big|_0^\infty - \int_0^\infty t \cdot F'(t)\,dt$$

$$= 0 + \int_0^\infty t \cdot \big(-F'(t)\big)\,dt$$

$$= \int_0^\infty t\,f(t)\,dt$$

$$= \langle T \rangle$$

**Supplementary Note 3. Commentary on the assumption that adsorption reaction is pseudo first-order**

A common model for phage adsorption assumes that the free phage depletion rate depends on the concentration of both bacteria $B$ and phages $P$ , with adsorption rate constant $k$ (2–4):

$$\frac{dP}{dt} = kBP \qquad \text{Equation 2}$$

Solving this equation with initial condition $P(t = 0) = P_0$ yields

$$k = -\frac{1}{Bt}\ln\left(\frac{P}{P_0}\right) \qquad \text{Equation 3}$$

A number of studies in literature report adsorption rate constant $k$ with units [mL $min^{-1}$], obtained through the above model. $B$ is assumed to be constant throughout the 10-minute experiment because $B$ is in large excess of $P$. It is assumed that a 1.4-fold difference in $B$ (assuming fast growth, 20-minute doubling time, which results in 1.4-fold change in concentration within 10 minutes) does not have a huge impact on the bacterial surfaces that the phages have available for attachment(5). However, if $B$ is included in the calculation of $k$, the measurement errors in $B$ contribute to the errors in $k$ (**Fig S6**).

Following the same argument, under our experimental condition where $B$ (~0.5-5×$10^8$ CFU/mL) is in large excess relative to $P$ (~1-5×$10^4$ PFU/mL), an assumption of pseudo first-order phage depletion is fair(6), where $B$ is assumed to be constant in the model, and hence $kB$ in Equation 2 is replaced by an effective adsorption rate constant $k_{eff}$ [$min^{-1}$], which yields

$$\frac{dP}{dt} = k_{eff}P \qquad \text{Equation 4}$$

This assumption has been experimentally and argumentatively validated by classic literature (5, 7) and discussed by recent literature on phage adsorption(4). Solving Equation 4, we get Equation 1 (main text):

$$P = P_0 \exp\left(-k_{eff}\, t\right)$$

We followed this assumption in order to get a less error-prone estimate of rate constant from the classic adsorption assay (see replicate distributions and error-bars in **Fig S6**A versus those in **Fig 3**C), which we could compare with the microscopy assay outcomes.

**Supplementary figures**

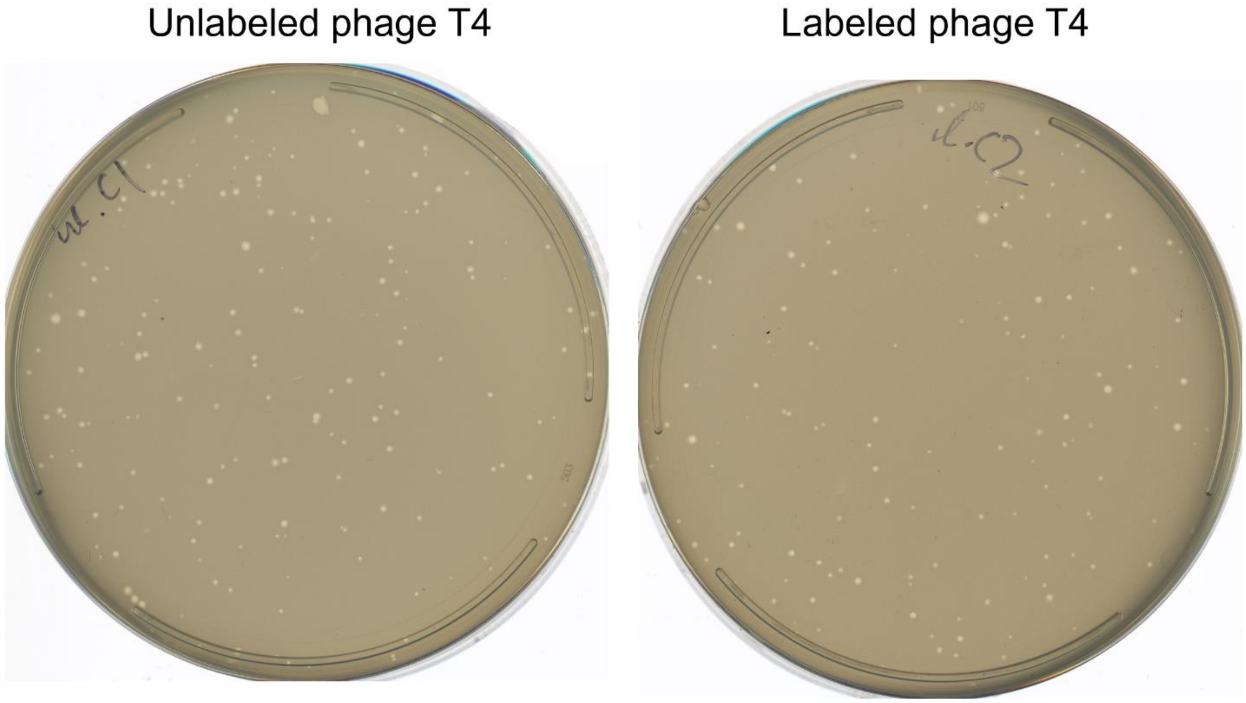


**Fig S1. Labeled phages retain their infectivity.** Plaque assays were performed with labeled and unlabeled phage T4 with host BW25113. Labeled phages yielded plaques on lawns of the bacterial hosts, indicating successful infections. We also performed similar experiments with all other phages used in this study (phages λ, T5, M13, T2, T7, P1, A1-1, and OMKO1) and observed identical results (data not shown).

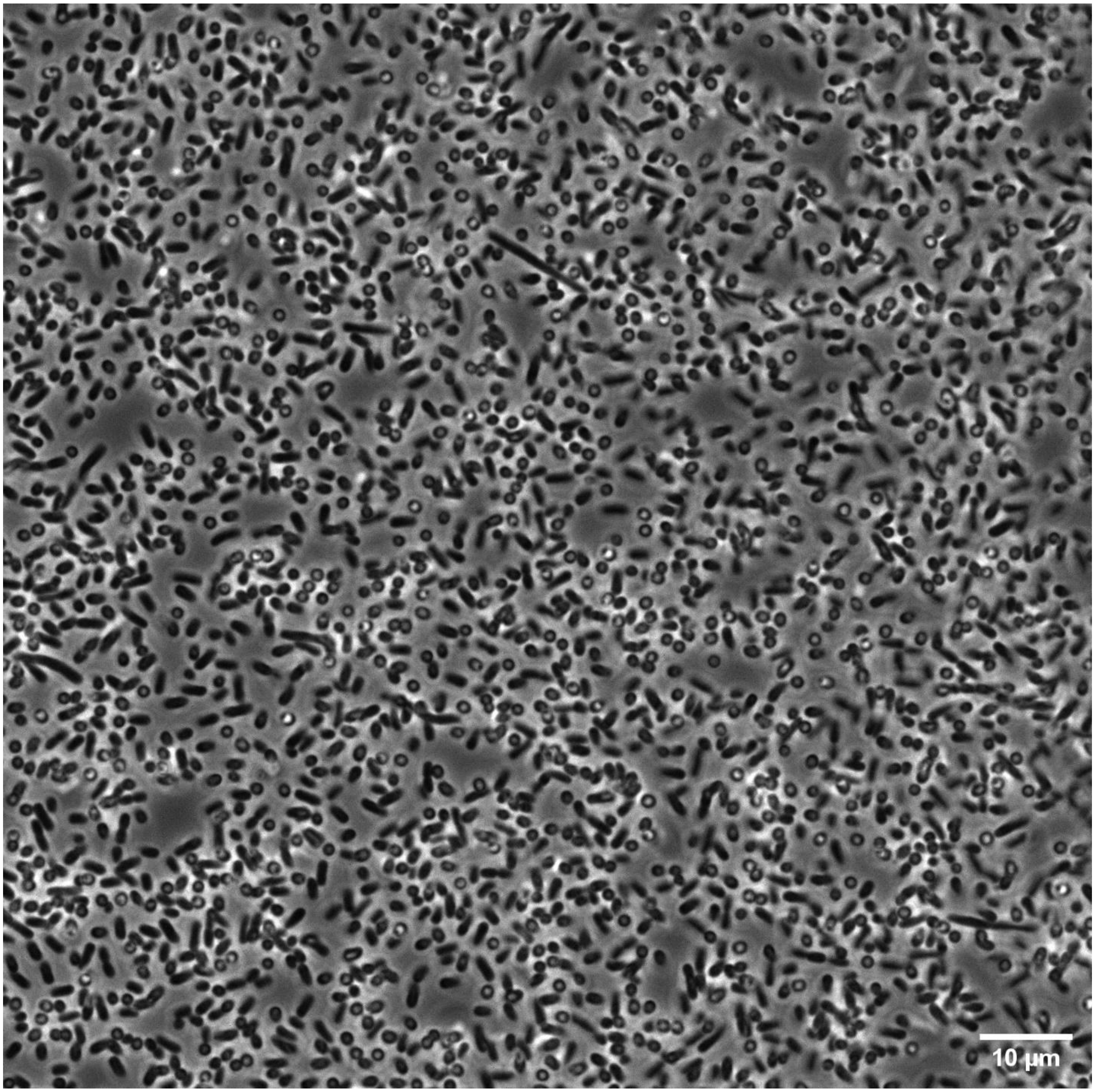


**Fig S2. Lawn of closely packed bacteria immobilized on the coverslip.** Phages were introduced to this lawn after immobilizing bacteria on the glass coverslip with 0.01% poly-L-lysine.

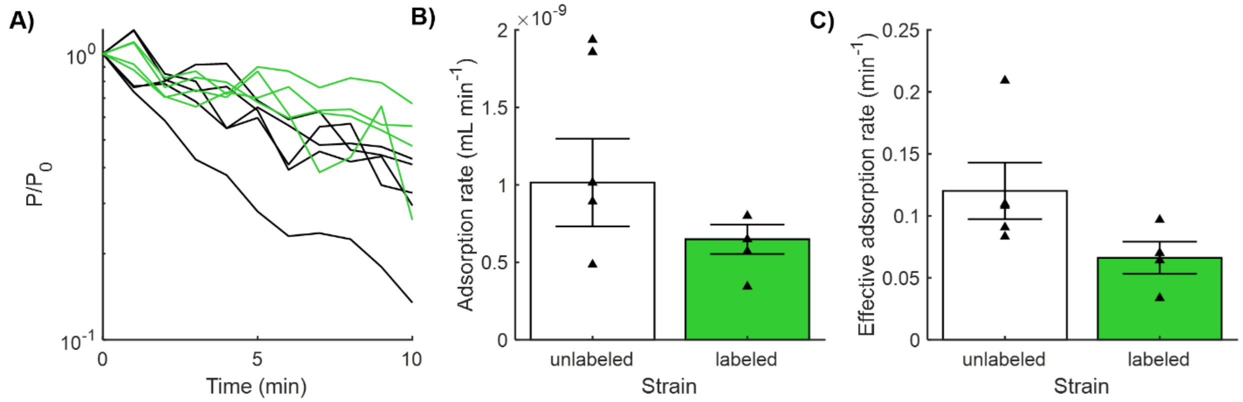


**Fig S3. Classical adsorption assay revealed that adsorption rates of labeled and unlabeled phages are comparable. (A)** Classic adsorption curves for T4 phages, unlabeled (black lines; n = 5 replicates) or labeled with the lysine-specific dye (green lines; n = 4 replicates) are indicated. Estimates of the adsorption rate constant $\boldsymbol{k}$ **(B)** and the effective adsorption rate constant $\boldsymbol{k_{eff}}$ **(C)** showed no statistically significant differences, when comparing labeled and unlabeled phages. Statistical significance was assessed by Student's $t$-test, which yielded P = 0.09 in each comparison. Plots depict means ± standard errors.

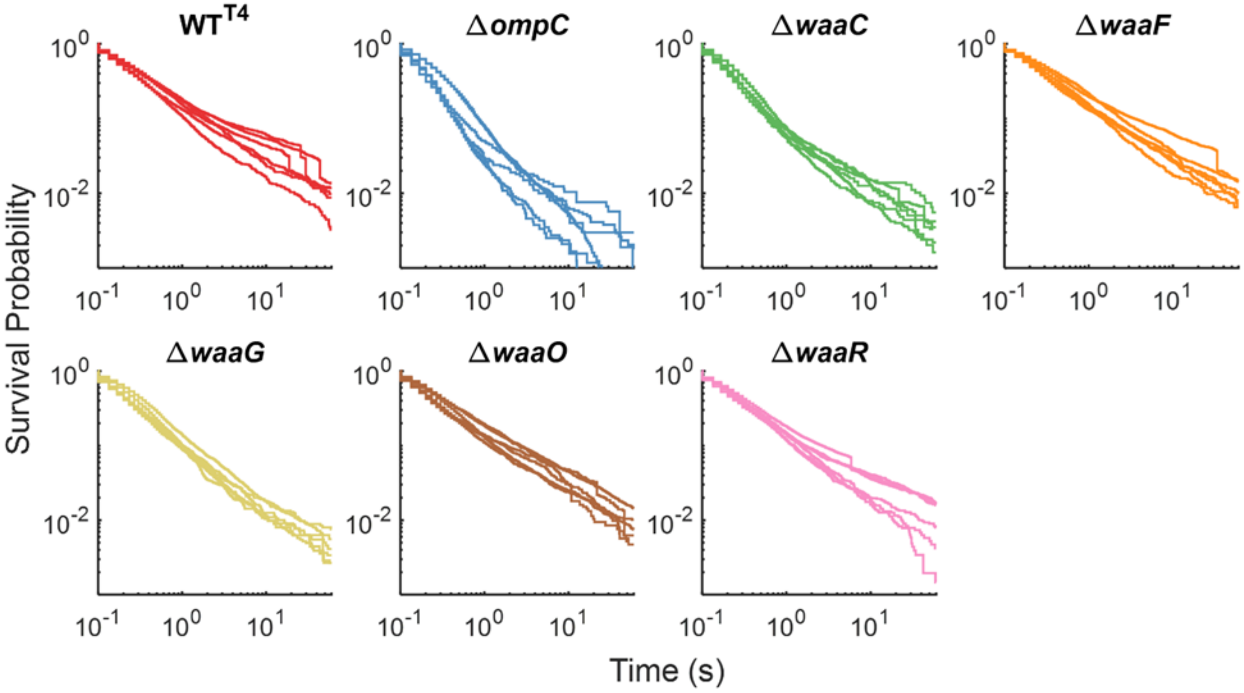


**Fig S4.** Survival probability curves (n = 6 replicates) for phage T4 on each bacterial strain shown in **Fig 2**D.

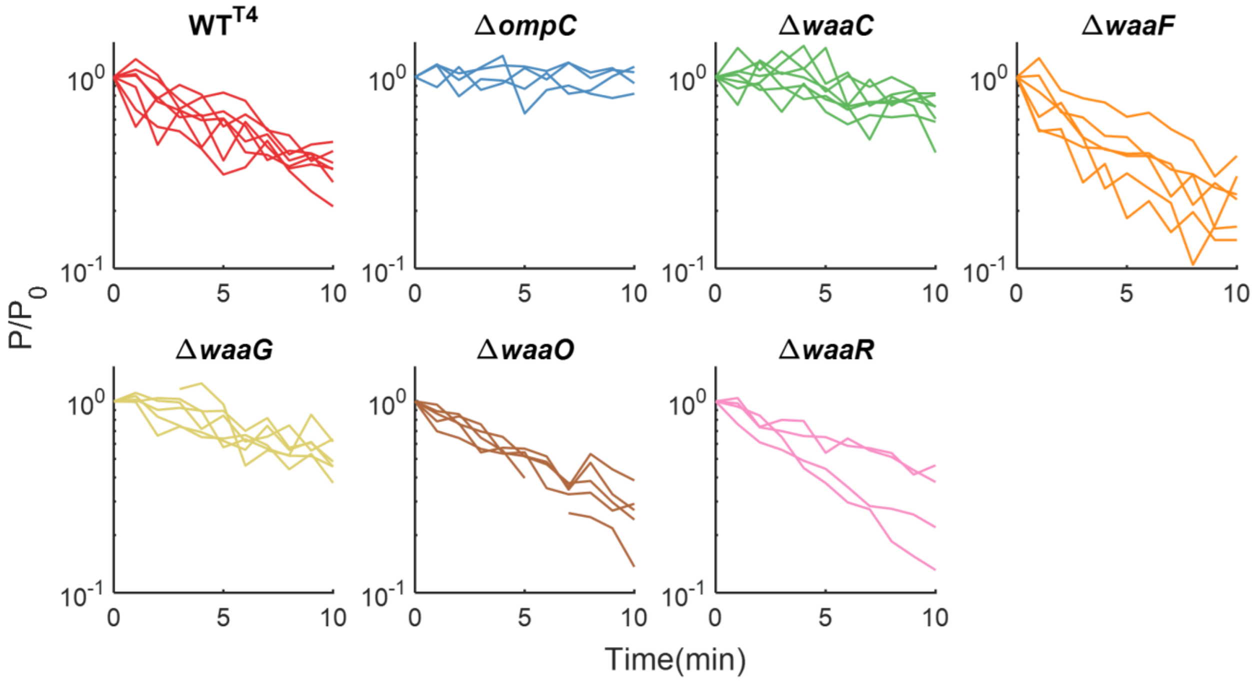


**Fig S5.** Classic adsorption curves (n = 4 to 6 replicates) for phage T4 on different host strains of *E. coli* shown in **Fig 3**C.

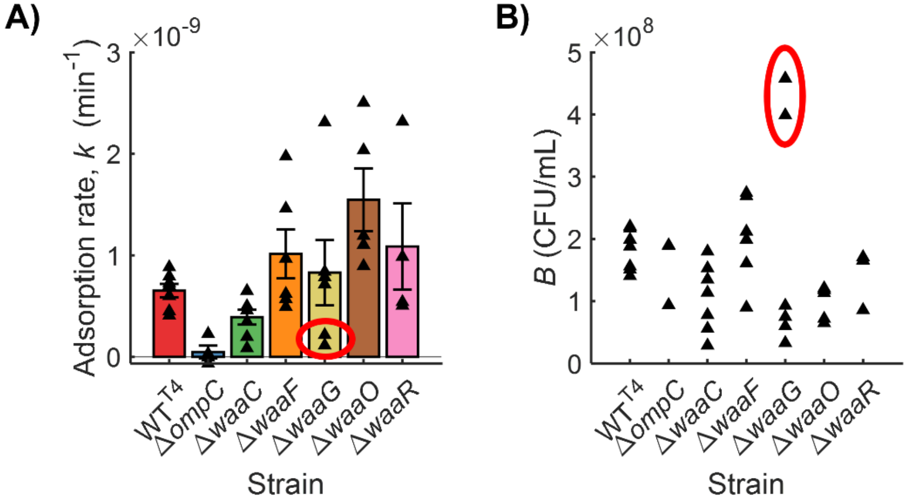


**Fig S6. Adsorption rate constant $k$ has a higher measurement variability due to measurement variability in bacterial concentration. (A)** Adsorption rate constant $k$ was calculated for each experimental replicate as $k = -\frac{1}{Bt}\ln\left(\frac{P}{P_0}\right)$, which accounts for the bacterial concentration $B$. **(B)** Bacterial concentration measured for each experimental replicate is plotted. The datapoints within the red circles exemplify how extreme values of $B$ in the experiment influence the calculated value of $k$.

**Movie S1** (separate file). After addition of fluorescently labeled phages, a majority of bacteria in the microscopic focal view visibly undergo lysis.
